## Supplementary Materials for "FinaleMe: Predicting DNA methylation by the fragmentation patterns of plasma cell-free DNA"

**Supplementary Figures**

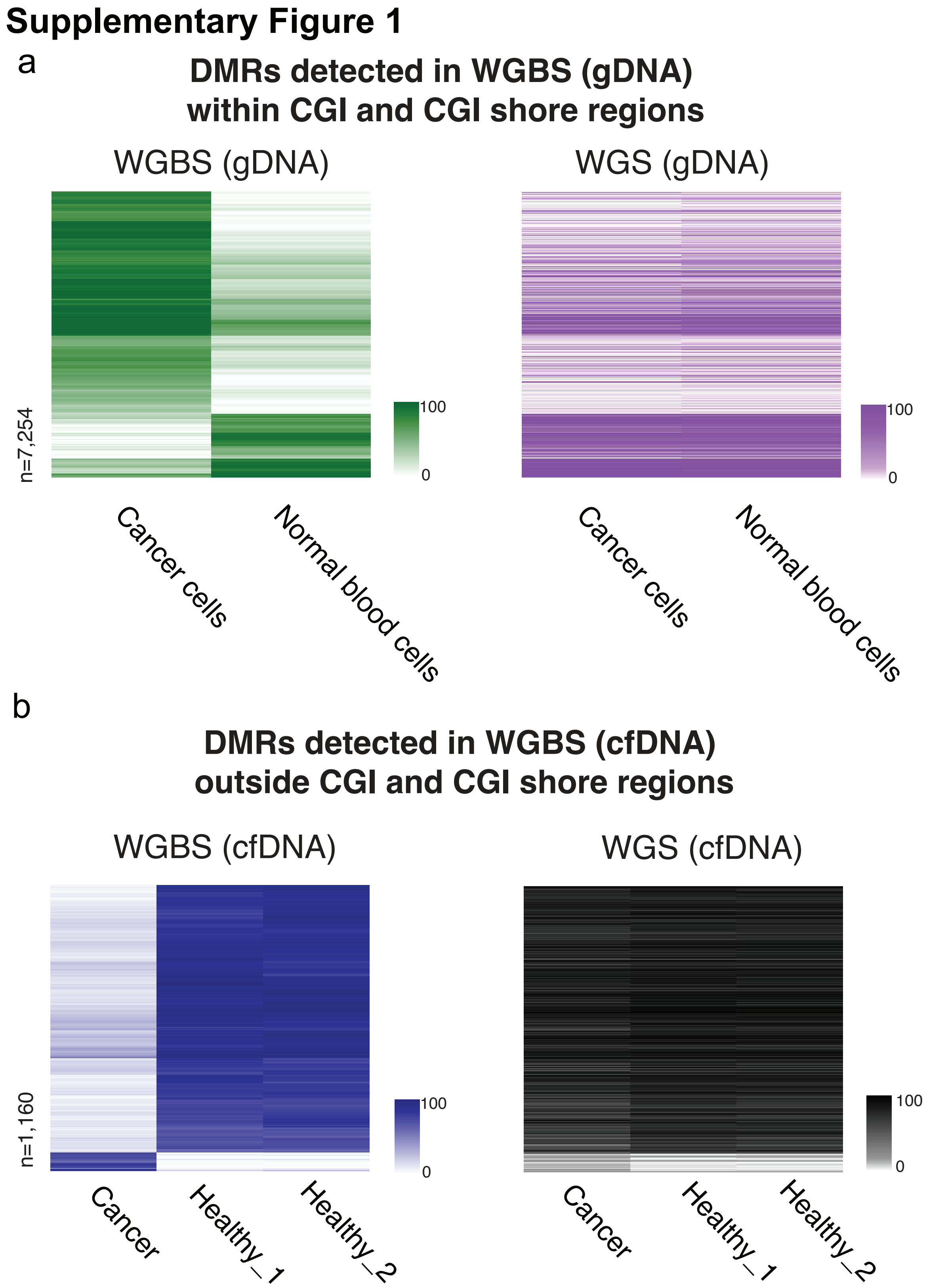


**Supplementary Figure 1**. Heatmap of measured (left panel, WGBS) and predicted (right panel, WGS) DNA methylation level at differentially methylated windows (1kb) characterized in WGBS. a. results in gDNA at

CGI and CGI shore regions. b. results in cfDNA at CpG-poor regions (no CGI or CGI shore regions in +/-2kb). The row orders in both WGBS and WGS datasets were based on the clustering of DNA methylation levels in WGBS only.

Supplementary Figure 2

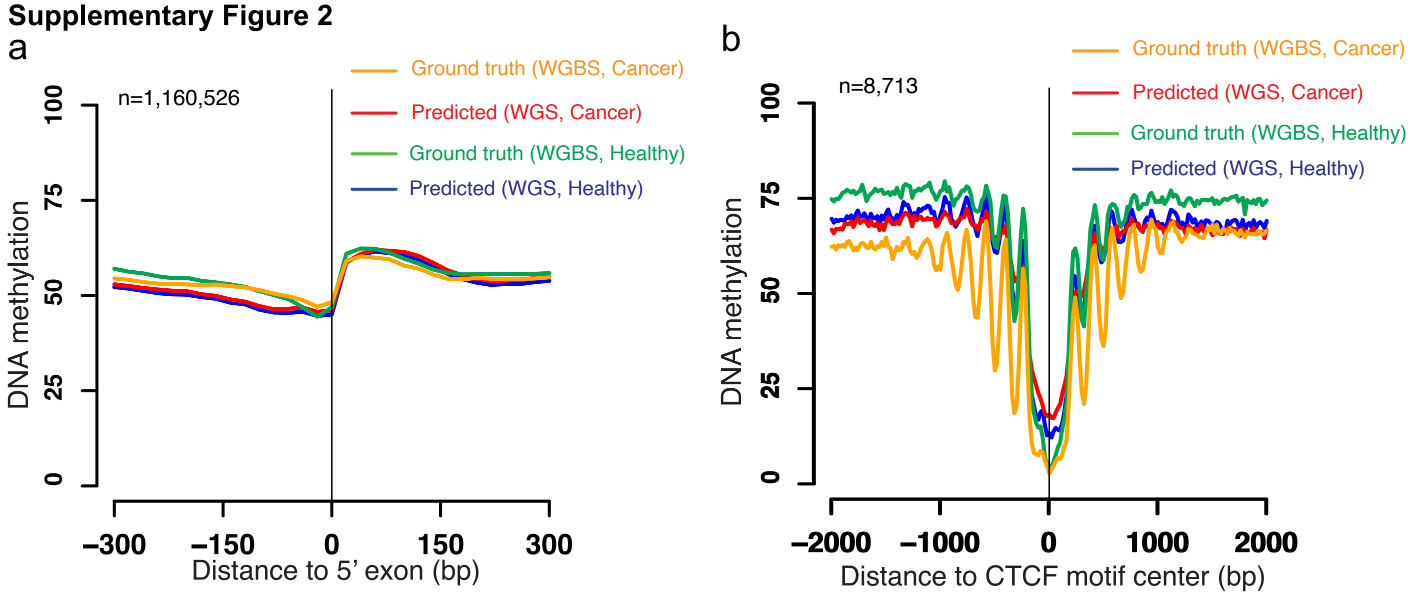


**Supplementary Figure 2**. Average ground truth (WGBS) and predicted (WGS) DNA methylation level at a. exons and b. CTCF motif from cancer and healthy individuals.

### Supplementary Figure 3

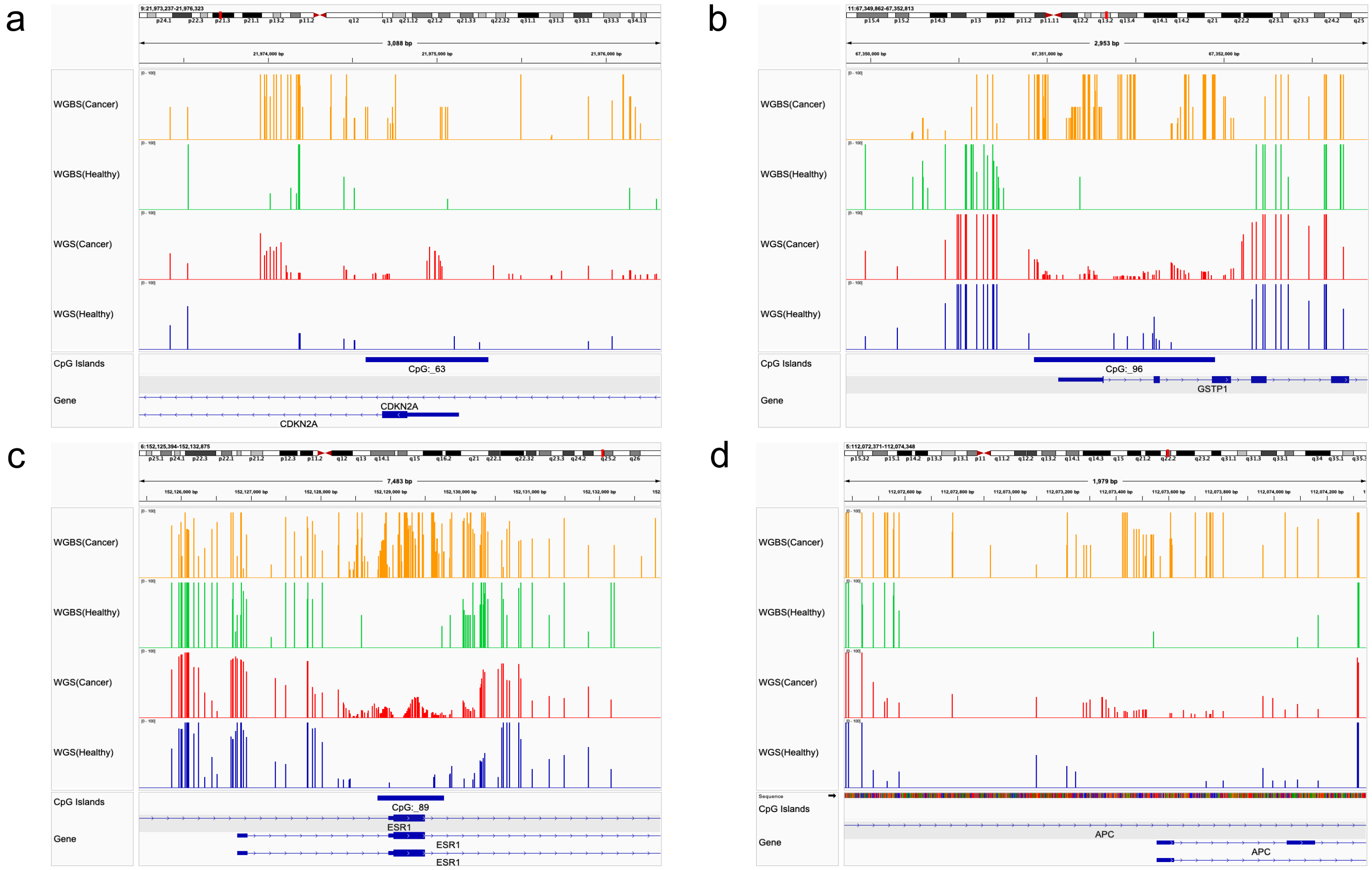


**Supplementary Figure 3**. Example regions that are often hypermethylated in prostate cancer patients. a. CDKN2A, b. GSTP1, c. ESR1, d. APC.

**Supplementary Figure 4**

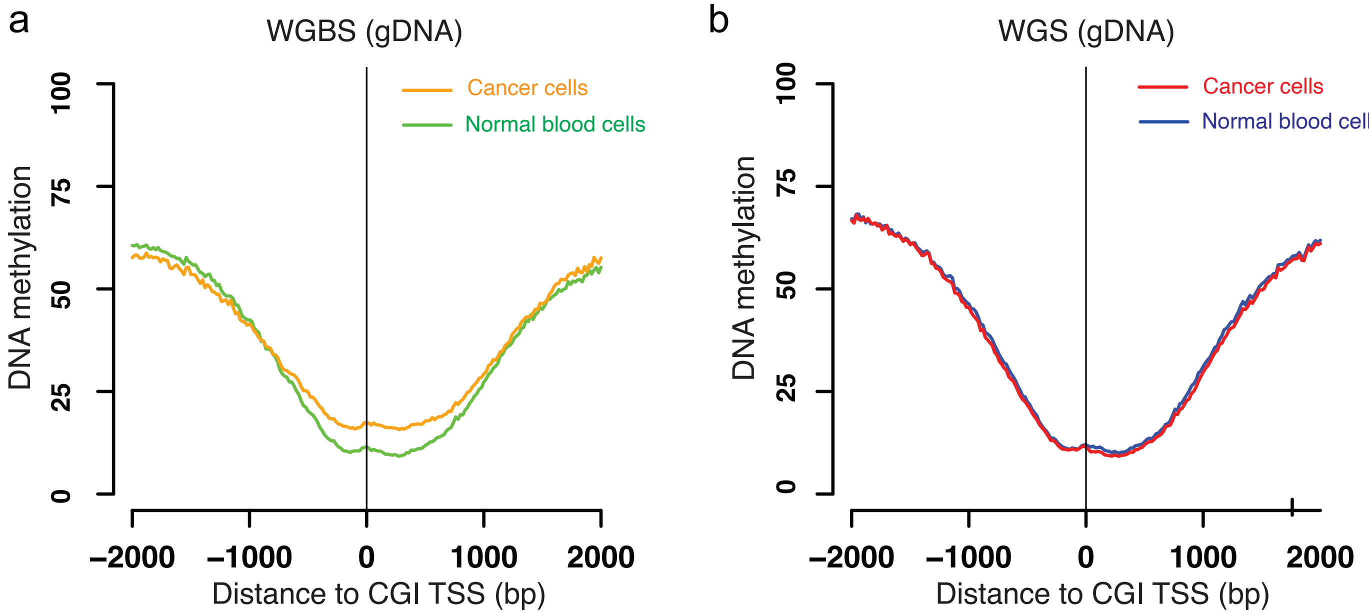


**Supplementary Figure 4**. Average DNA methylation level at CpG island promoter region from gDNA obtained in cancer cells (HepG2, liver cancer cell line) and normal blood cells (GM12878, B-lymphoblastoid cell line) at a. ground truth (WGBS) and b. predicted (WGS).

### Supplementary Figure 5

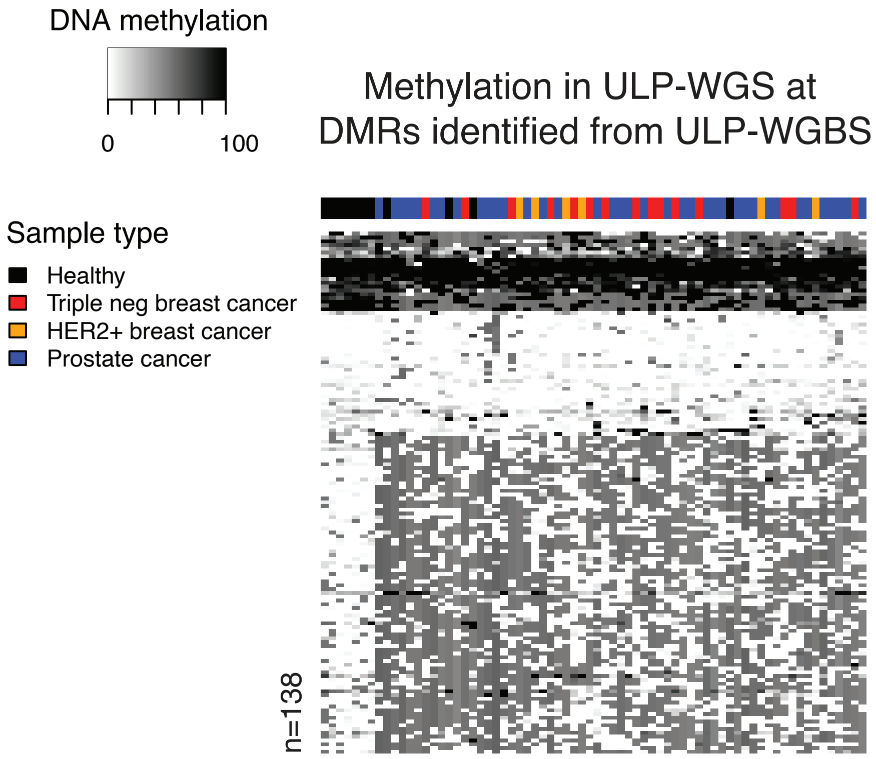


**Supplementary Figure 5**. Heatmap of predicted DNA methylation level in ULP-WGS at differentially methylated windows (1kb) characterized in ULP-WGBS between cancers and healthy individuals.

**Supplementary Figure 6**

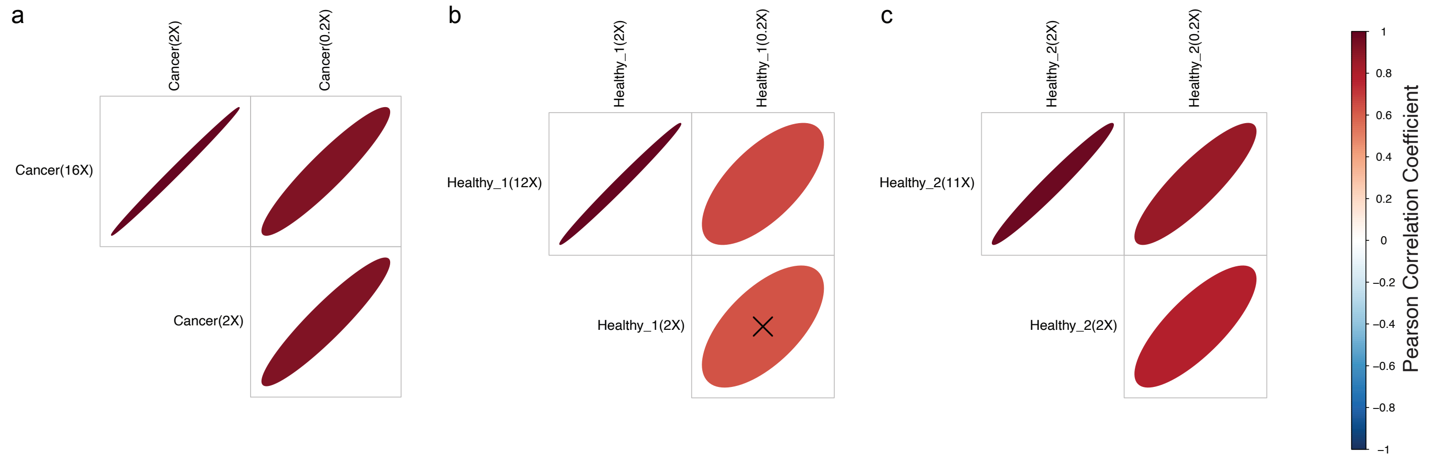


**Supplementary Figure 6**. The correlation of tissues-of-origin predictions results between deep cfDNA WGBS and their downsampled WGBS dataset from a. cancer, b. healthy (HD_45), and c. healthy (HD_46) individuals. The percentage of tissues that contributed to cfDNA was first calculated in each sample. Then the correlation between these tissues-of-origin vectors was calculated and compared between high-coverage ones and downsampled low-coverage ones. "corrplot" package in R was utilized to visualize the correlation. "X" on top of the plot means that "the correlation is not statistically significant ($p>0.05$)". The shape of the plot represents the dispersion status of the dot.

Supplementary Figure 7

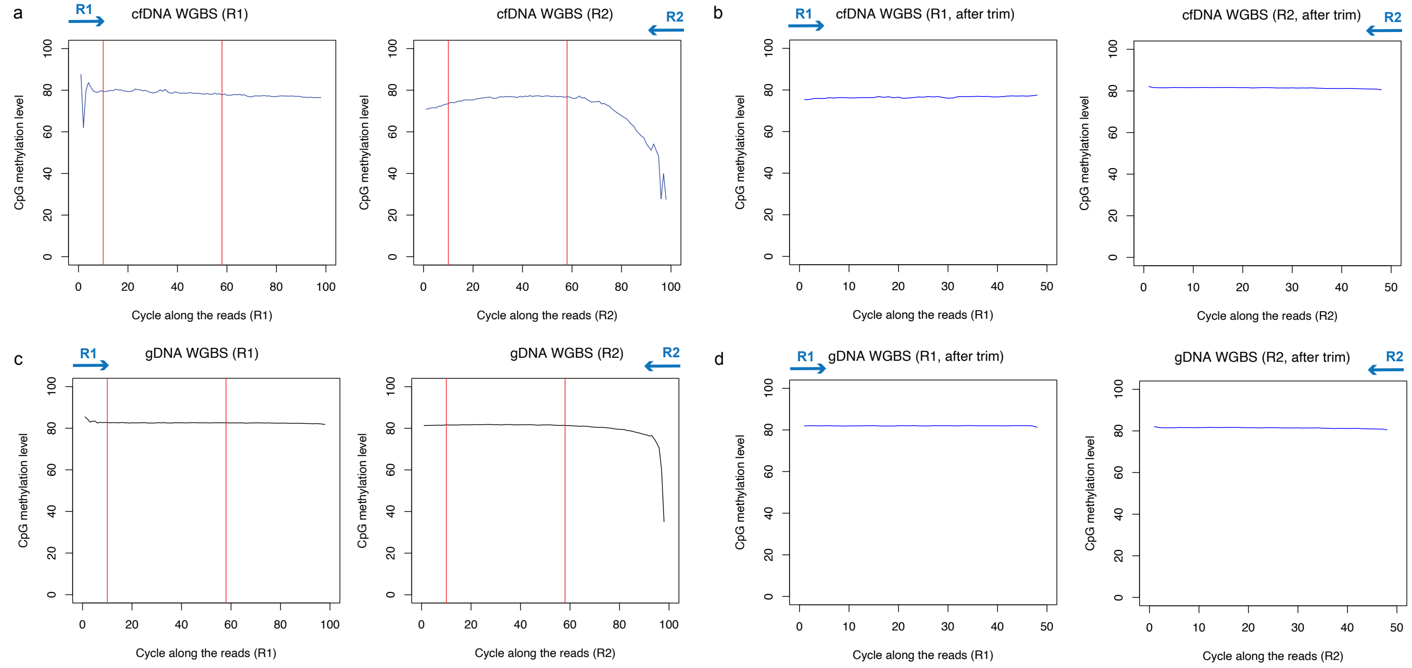


**Supplementary Figure 7**. M-bias plot to characterize the part of reads that are potentially affected by jagged-end in cfDNA WGBS. The analysis was done at both a. cfDNA and b. gDNA. Red lines were the cut-off used to trim the reads in WGBS.

Supplementary Figure 8

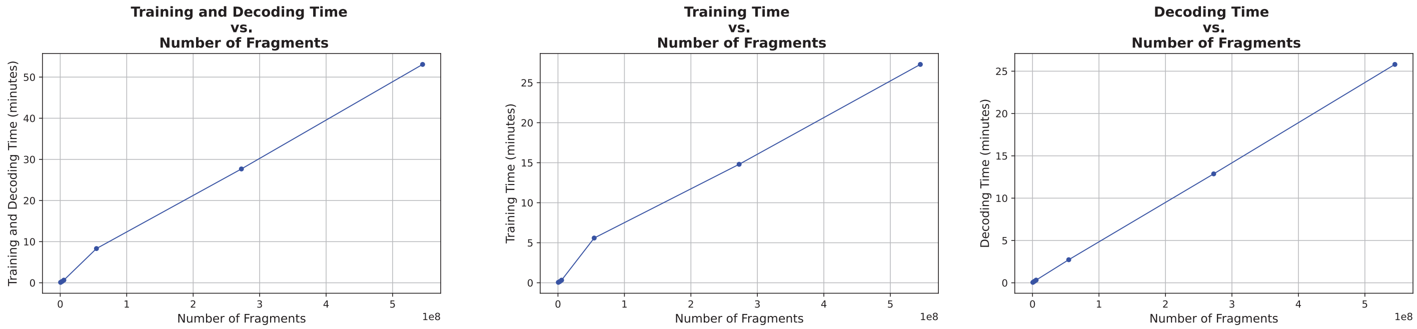


**Supplementary Figure 8**. The model's time cost (minutes) on the cfDNA WGS dataset (1 million-600 million fragments). Benchmark was performed at a single CPU in the computational cluster (Intel(R) Xeon(R) Gold 6338 CPU @ 2.0GHz).

**Supplementary Table 1**. Meta-data information for each sample.

**Supplementary Table 2.** Batch information for cfDNA WGS data.

**Supplementary Table 3**. Batch information for cfDNA WGBS data
